# Multi-trait multi-environment genomic prediction of preliminary yield trials in pulse crops

**DOI:** 10.1101/2024.02.18.580909

**Authors:** Rica Amor Saludares, Sikiru Adeniyi Atanda, Lisa Piche, Hannah Worral, Francoise Dariva, Kevin McPhee, Nonoy Bandillo

## Abstract

Phenotypic selection in preliminary yield trials (PYT) is challenged by limited seeds, resulting in trials with few replications and environments. The emergence of multi-trait multi-environment enabled genomic prediction (MTME-GP) offers opportunity for enhancing prediction accuracy and genetic gain across multiple traits and diverse environments. Using a set of 300 advanced breeding lines in the North Dakota State University (NDSU) pulse crop breeding program, we assessed the efficiency of a MTME-GP model for improving seed yield and protein content in field peas in stress and non-stress environments. MTME-GP significantly improved predictive ability, improving up to 2.5-fold, particularly when a significant number of genotypes overlapped across environments. Heritability of the training environments contributed significantly to the overall prediction of the model. Average predictive ability ranged from 3 to 7-folds when environments with low heritability were excluded from the training set. Overall, the Reproducing Kernel Hilbert Spaces (RKHS) model consistently resulted in improved predictive ability across all breeding scenarios considered in our study. Our results lay the groundwork for further exploration, including integration of diverse traits, incorporation of deep learning techniques, and the utilization of multi-omics data in predictive modeling.

**Core ideas:**

- Phenotypic selection in PYT is challenged by limited seeds, resulting to few replications and environments.
- MTME-GP offers opportunity for enhancing prediction accuracy of multi-trait and diverse environments in PYT.
- MTME-GP enhances prediction by up to 2.5-fold, especially with numerous overlapping genotypes in various tested environments.
- RKHS MTME-GP models, excels in low-heritability, negatively correlated traits, like drought-affected conditions.

## 1.0 INTRODUCTION

The challenges posed by a rapidly expanding global population and climate change underscore the imperative for sustainable food production (Tilman et al. 2011; van Dijk et al. 2021; Kumar et al. 2022). Field pea (*Pisum sativum*) emerges as a desirable crop, not only meeting the criteria for sustainability but also standing out as an affordable and nutritious plant-based protein source. This places field pea at the forefront of leguminous crops in the food industry (Punia & Kumar, 2022; Shanthakumar et al., 2022). However, the conventional process of developing a promising line for release to farmers involves rigorous phenotypic assessments across multiple seasons and environments, especially for polygenic traits with complex genetic architecture (Samantara et al., 2022). Accelerating the development of crop varieties to meet the needs of a growing population stands out as a viable strategy to help feed the world (Ahmar et al. 2020).

Genomic selection for complex traits in early breeding cycles has the potential to significantly reduce the selection cycle time and expedite genetic gain (Ertiro et al., 2015; Crossa et al., 2017; Bernardo, 2020). The advent of next-generation sequencing and various genotyping platforms has rendered genotyping more accessible and cost-effective than traditional phenotyping methods (Atanda et al., 2021). This transformative shift provides a unique opportunity to seamlessly integrate genomic selection (GS), leveraging DNA information to predict the genetic merit of new genotypes (Meuwissen et al., 2001; Atanda et al., 2021). Studies have shown the potential of GS in pulse breeding programs for genetic improvement of seed yield, seed protein content, and wider adaptability to ever-changing environmental conditions (Annicchiarico et al., 2019; Budhlakoti et al., 2022; Cazzola et al., 2021; Gosal & Wani, 2020; Haile et al., 2020; Li et al., 2022; Pratap et al., 2022). The North Dakota State University (NDSU) pulse breeding program is undergoing a fundamental shift from phenotypically-driven approaches to a more modern GS-based approach at the preliminary yield trial (PYT) stage. Improving accuracy in the early yield testing stage for selection of top-performing lines is essential for efficient resource allocation, shortening the breeding cycle, and, ultimately, increasing genetic gain (Bassi et al., 2016; Atanda et al., 2021; Bandillo et al., 2022).

Univariate or single-trait (UNI) models have been widely employed in GS, focusing on predicting individual traits independently while assuming no correlation between traits (Atanda et al., 2022; Sandhu et al., 2022; Montesinos-López et al., 2022). Multi-trait GS (MT-GS) models integrate information from correlated traits and shared genetic information between lines to improve the accuracy. (Jia and Jannink, 2012; Gill et al. 2021; Atanda et al., 2022; Montesinos-López et al., 2022;). As traits are genetically correlated, these MT-GS models have demonstrated their ability to enhance prediction accuracy, particularly for traits with inherently low heritability.

Hayes et al. (2017) reported increased genomic prediction accuracy by ∼40% for wheat end-use quality traits using a MT-GS model compared to a UNI-GS model. In barley, Bhatta et al. (2020) reported an increase of 57 to 61% prediction accuracy for agronomic and malting quality traits. In a recent study, Atanda et al. (2022) proposed a sparse-phenotyping-aided MT-GS model and demonstrated a notable improvement of over 12% in prediction accuracy across nutritional traits in field pea. Generally, prediction accuracy in MT-GS improves as correlation between traits increases. However, in practice, the correlation between traits ranges from positive to negative, along with varying degrees of heritability. Addressing this challenge, Atanda et al. (2022) emphasized composition of traits in the training and prediction sets based on the heritability and genetic correlation between traits to enhance the prediction accuracy. Studies have also shown that the integration of genotype by environment (GxE) in the MT model further improves prediction accuracy (Gill et al., 2021; Sandhu et al., 2022).

In this study, we explored the merit of a multi-trait multi-environment enabled genomic prediction model (MTME-GP) in enhancing the prediction accuracy of two highly-important, yet negatively correlated, traits: seed protein content and seed yield in field pea. Additionally, we further assessed the potential of MTME-GP models for predicting performance within- and across-environments using multiple years of data.

## 2.0 MATERIALS AND METHODS

### 2.1 Germplasm and phenotyping

The genetic materials consisted of 282 NDSU advanced elite breeding lines previously described in Bari et al. (2022). The lines were planted in 1.5-x 7.6-m plots at 0.30-m spacing between plots with 840 pure live seeds per plot, arranged in an augmented incomplete block design with five diagonal repeated checks for preliminary yield trials. Seed yield and agronomic data were collected in 3-year experiments from 2020 to 2022, including two environments at the NDSU North Central Research Extension Center (NCREC) near Minot, ND (MOT20 and MOT21) and one environment at the Carrington Research Extension Center near Carrington, ND (CAR22). Standard cultural practices were followed. Plots were harvested at physiological maturity (90-120 days after planting) and dried to 13% moisture content. A total of 0.11 kg clean and dried harvested seeds per line was used for protein analysis at the NCREC using near infrared (NIR) spectroscopy.

### 2.2 Genotyping

Young leaves were harvested from seedlings of each pea line planted in a greenhouse environment. DNA extraction was carried out using the DNeasy® Plant Mini Kit (Qiagen, Germantown, MD, USA) following the manufacturer’s instructions, and elution was performed with 100µl. Subsequently, the DNA samples obtained were quantified using the Qubit dsDNA BR Assay kit and Qubit 4.0 fluorometer (Life Technologies Corporation, Eugene, OR). As described by Bari et al. (2022), DNA samples were standardized to a final concentration of 25 ng/µl for subsequent genotyping-by-sequencing (GBS) at a genomic center. The prepared dual-indexed GBS libraries using the restriction enzyme *ApeKI* (Elshire et al. 2011) were combined into a single pool and sequenced across 1.5 lanes of NovaSeq S1x100-pb run, producing approximately 1,000 million pass filter reads with mean quality scores of > 30. The resulting quality reads were aligned to the established pea reference genome (Kreplak et al. 2019) yielding a total of 28,832 SNP markers. After removal of SNPs with minor allele frequency less than 1%, heterozygosity exceeding 20%, and those having over 90% missing values, the remaining 11,858 SNPs were used for downstream analysis. SNPs with missing values were imputed using Beagle v.5.1 (Browning et al., 2018).

### 2.3 Phenotyping

A mixed linear model was used to extract best linear unbiased estimates (BLUEs) for all traits evaluated using the following model:

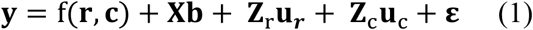

where **y** is the response variable for n-th phenotype, **b** is the fixed effect of the genotype, **u**_**r**_ and **u**_**c**_ are row and column random effects accounting for discontinuous field variation with multivariate normal distribution: 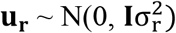 and 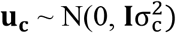 respectively, wherein, **I** is an identity matrix and 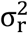 and 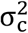 are variances due to row and column effect. f(**r, c**) is a smooth bivariate function defined over the row and column positions, **ε** is the measurement error from each plot with distribution of 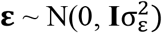, wherein, **I** is the same as above and 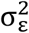 is variance for the residual term or simply referred to as nugget. **X** and **Z** are incidence matrices for the fixed and random terms, respectively.

### 2.4 Genomic selection models

The univariate (UNI) single environment GS model was fitted using the Bayesian approach and implemented in the BGLR R package (Pérez & de los Campos, 2014):

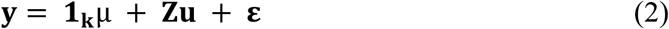

where **y** is the vector (n x 1) of adjusted means (BLUEs) for j-th pea lines for a targeted trait; μ is the overall mean; **1**_k_ (k × 1) is a vector of ones; **u** is the genomic effect of the j-th pea line and assumed to follow the multivariate normal distribution expressed as 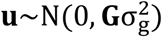, where **G**is the genomic relationship matrix and 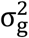 is the additive genetic variance; and **Z** is the incidence matrix for genomic effect of the lines.

The UNI multi-environment GS model was fitted using a reaction norm model which accounts for genotype by environment interaction (GxE) described in Jarquin et al. (2013):

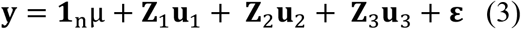

where **y** (n×1) is the vector of phenotypes of the pea lines measured in the environments (1…k), μ is the overall mean and **1**_n_ (nx1) is a of vector ones. **u**_1_is the random effect of the k-th environment and follows the multivariate normal distribution 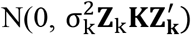 where 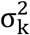 is the variance of the main effect of the environment, **K** is a relationship matrix between the environments which is an identity matrix, **Z**_k_ is an incidence matrix that relates the phenotypes to the mean of the environments, and 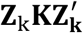 is a block diagonal matrix that uses a 1 for all pairs of observations in the same environment and a 0 for off-diagonal elements. **u**_**2**_is the random effect of the pea lines and follows the multivariate normal distribution 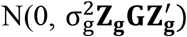, where 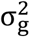 is the variance of the main effect of the pea lines, **Z**_g_is an incidence matrix that relates the phenotypes with the genomic relationship between the pea lines (**G). u**_3_is the random effect of the GxE effect and follows the multivariate normal distribution 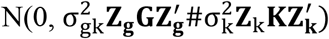, where 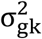 is the variance component of GE, # denotes the Hadamard product, and 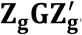 and 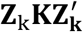 are the same as previously described. **ε** is the random term of the residual and follows the multivariate normal distribution 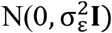, where 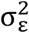 is the homogenous residual variance. For the Bayesian Reproducing Kernel Hilbert Spaces Regressions (RHKS), the **G** matrix was replaced by kernel matrix (see Pérez & de los Campos, 2014 for details).

The multi-trait (MT) single environment GS model was fitted by extending Eq. 2 as follows:

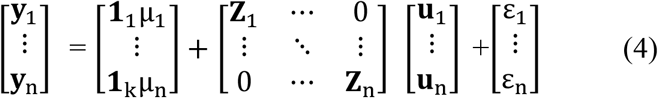

where **y**_1_…**y**_n_ are the vector of phenotypes, μ_1_ …μ_n_ are the overall mean for each n-th trait, **Z**_1_ …**Z**_n_ is the incidence matrix for genomic effect of the lines for each n-th trait, **u**_1_ …. **u**_n_ is the genomic effect of the lines for each n-th trait, and **ε**_1_ … **ε**_n_ is the residual error for each n-th trait. The random term is assumed to follow the multivariate normal distribution [**u**_1_ …. **u**_n_] ∼ MN[0, (**G**⨂ **G**_o_)], where **G** is the same as above and **G**_o_ is an n x n unstructured variance-covariance matrix of the genetic effect of the traits, this is represented as follows:

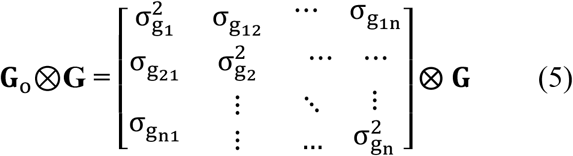

The diagonal elements represent variance for each trait and covariances between traits are the off-diagonal elements.

Further, the residual term for each n-th trait is assumed to follow the multivariate normal distribution [**ε**_1_ … **ε**_n_] ∼ MN[0, (**I** ⨂ **R**)], where **I** is the same as above and **R** is a heterogeneous diagonal matrix of the residual variances for each n-th trait:

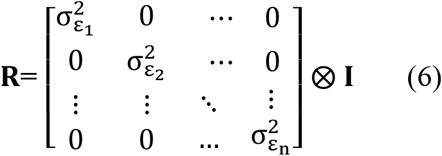

The diagonal elements represent the residual variance for each n-th trait and off-diagonal elements of the **R** matrix equal zero.

For the multi-trait (MT) multi-environment GS model, Eq. 3 was expanded as described by Montesinos et al. (2022):

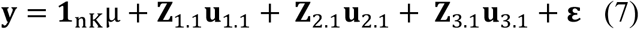

where **y** is of size i x n and i =j x k, n is the number of traits, j is the number of genotypes and k is the number of environments. **Z**_1.1_ is the incidence matrix of environment of size i x k, **u**_1.1_ is the random effect of each environment of each trait with size k x n, **Z**_2_.1 is the incidence matrix of genotypes of order i × j, **u**_2.1_ is the random effect of the genotypes i × n, and follows the multivariate normal distribution 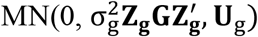, where **Z**_**g**_ is an incidence matrix of the genotypes of order i x j. 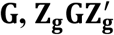 and 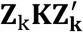 are the same as above and **Ug** is the unstructured variance-covariance matrix of traits of order n × n. **Z**_3.1_ is the incidence matrix of GE of order i × kj, **u**_3.1_ is the random effect of the genotypes by environment by trait of order kj × n and follows the matrix multivariate normal distribution MN 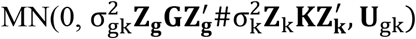, where **U**_gk_ is the unstructured variance-covariance matrix of order k by k. **ε** is the random term of the residual and follows the multivariate normal distribution MN(0, **I**, Σ_t_). **I** is identity matrix of order i ×n, and Σ_t_ is the unstructured variance-covariance matrix.

### 2.5 Cross validation scheme

Evaluation of the predictive performance was assessed using various validation scenarios mean to mimic possible utilization scenarios of genomic selection in the NDSU field pea breeding program. Models were trained to predict seed yield and total seed protein content within and across different environments. Predictive ability (PA) was estimated as the Pearson correlation coefficient between predicted genomic estimated breeding value (GEBV) and BLUE of each trait for the complete dataset. For within-environment predictions, datasets for each environment (MOT20, MOT21, and CAR22) were partitioned into different training set sizes (50%, 60%, 70%, and 80%) and the process was repeated 30 times. For across-environment predictions, models were trained on one environment and tested on a novel environment (e.g., MOT20 trained to predict MOT21). We also explored the effectiveness of training the model on multiple environments to predict a novel environment (e.g., MOT20 and MOT21 trained to predict CAR22).

## 3.0 RESULTS AND DISCUSSION

### 3.1 Predictive ability of different genomic prediction models

We evaluated the potential of GS to predict the genetic merit of two negatively-correlated complex traits across three environments with varying degrees of heritability (**Supp. Fig. 1**). Except in the case of UNI-GS model where G_BRR outperformed RKHS for both traits, the RKHS model consistently demonstrated superior predictive performance across models and in cross-validation strategies for both traits (**Supp. Fig.2:5**). The superiority of the RKHS model in all other scenarios evaluated in this study suggests robustness and reliability of the model in capturing not only additive effects but also non-linear effects and complex GxE interactions (Baertschi et al. 2021; Jiang and Reif 2015). These findings align with those of Bari et al. (2021), which observed subtle but favorable advantages of the RKHS model for predicting seed yield in field peas. To compare UNI with MT, we focused on the RKHS model due to its superiority over the BRR model across all validation scenarios. MT outperformed UNI by 11-fold across traits and environments (**Supp. Fig 6**:**7**). Okeke et al. (2017) also reported an improvement in predictive ability (average of 40%) with MT compared to UN for various traits in African cassava. Similarly, Arojju et al. (2020) reported improvements in prediction accuracy ranging from 24% to 59% for dry matter yield and 67% to 105% for nutritive quality traits in perennial ryegrass. Most recently, Winn et al. (2023) demonstrated substantial enhancement in prediction accuracy for various combinations of soft red winter wheat traits. These highlight the potential of MT models to enhance prediction accuracy, especially for challenging and resource-intensive traits.

The integration of GxE interaction by including environments with low heritability did not result in an increase in predictive ability as shown in Supplementary Figure 7F. Specifically, adding MOT21 to the training set, which has the lowest heritability estimate of 0.11, did not enhance predictive ability. This might suggest a nuanced interplay between environmental conditions and predictive models. This corroborates the findings of Rogers and Holland (2022), emphasizing that the addition of nuisance environments will not enhance overall predictive ability. This suggests that GxE interactions might be more relevant in environments with moderate to high heritability.

### 3.2 Optimal training set size for improved predictive performance of RKHS model

The training set size is one of the major factors influencing the prediction accuracy of un-tested lines (Norman et al., 2018). **Supp. Fig. 6:7** showed no clear trend in predictive ability for the traits with increasing training set size. The G_RKHS (MT) model consistently showed the highest predictive ability for seed protein reaching 30% when 60% and 80% of MOT20 dataset were trained to predict CAR22 (**Fig. 1**). However, in predicting for seed yield, the majority of the highest predictive abilities were still observed under G_RKHS (MT), reaching 34% when 80% of MOT20 dataset was trained to test CAR22 (**Supp. Fig. 8**). Previous studies have emphasized a strong relationship between prediction accuracy, training set size, and trait heritability (Luan et al., 2009; Lorenz et al., 2011; Clark et al., 2012; Nyline et al., 2017; Kaler et al.,2022; Atanda et al., 2022). Considering the varying heritability of our traits across environments, ranging from 1.57E-06 to 0.80 for grain yield and 0.12 to 0.53 for protein, and the negative correlation between traits, these factors might contribute to the overall predictive ability across models in our study. Contrary to our results, Bari et al. (2021) reported an increase in prediction accuracy with increased training set size. Other studies (Budhlakoti et al. 2022; De Roos et al. 2020) have also reported the influence of training set size and heritability on prediction accuracy. This underscores the importance of careful consideration when selecting training set size for model training.

**Figure 1.**
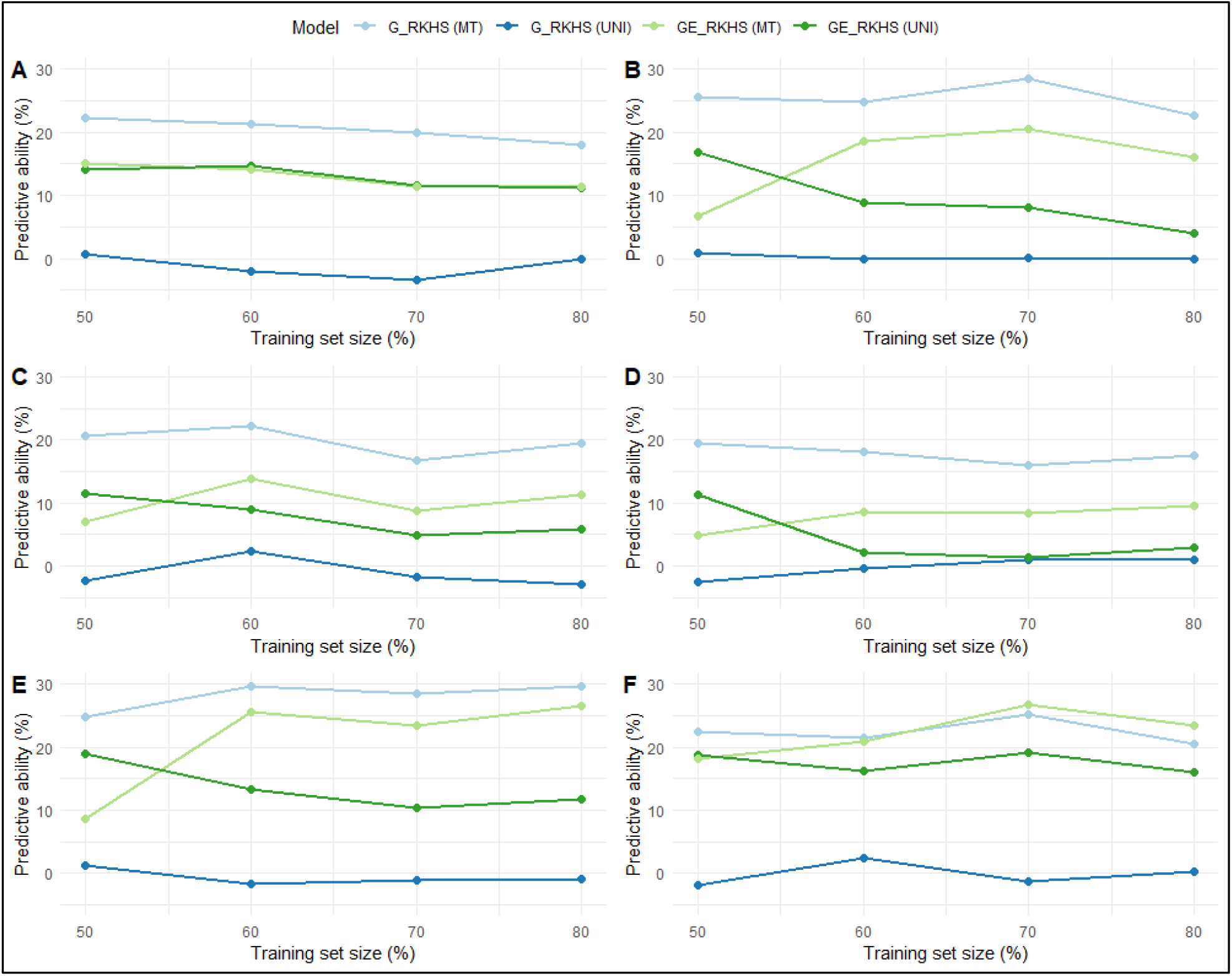
Average predictive ability with increasing training population size using RKHS models for seed protein content, RKHS is Reproducing Kernel Hilbert Spaces, MT is multivariate, UNI is univariate, G is prediction model considering genotype, GE is prediction model integrating GxE interaction. (A) MOT21 dataset trained to predict MOT20, (B) CAR22 dataset trained to predict MOT20, (C) MOT20 dataset trained to predict MOT21, (D) CAR22 dataset trained to predict MOT21, (E) MOT20 dataset trained to predict CAR22, (F) MOT21 dataset trained to predict CAR22.

### 3.3 Efficacy of MTME-GP for predictions within and across different environments

Generally, integration of GxE in the model improved the predictive ability (**Supp. Fig. 2B, 2E, 4B, 4E**) except for environment with low heritability as reported earlier. This further highlights the importance of carefully managing testing environments to reduce the influence of environmental nuisance on phenotyping. Ultimately, it underscores the significance of considering heritability in the environment when developing training datasets for multi-environment GS models, ensuring efficient capturing of the genetic relationship between environments and borrowing information effectively across environments (Xu, 2016; van Eeuwijk et al. 2019; Atanda et al., 2021). Similarly, Sapkota et al. (2020) reported varying prediction accuracy when environments with different heritability were included in the training model to predict new environments. Gill et al. (2021) emphasized the potential of MTME-GP in practical scenarios, such as overcoming the challenges posed by the loss of complete or partial trials due to extreme weather. As also shown in our study, also MTME-GP proved valuable in predicting the genetic merit of the lines affected by drought condition for both traits (**Fig. 2:3**).

**Figure 2.**
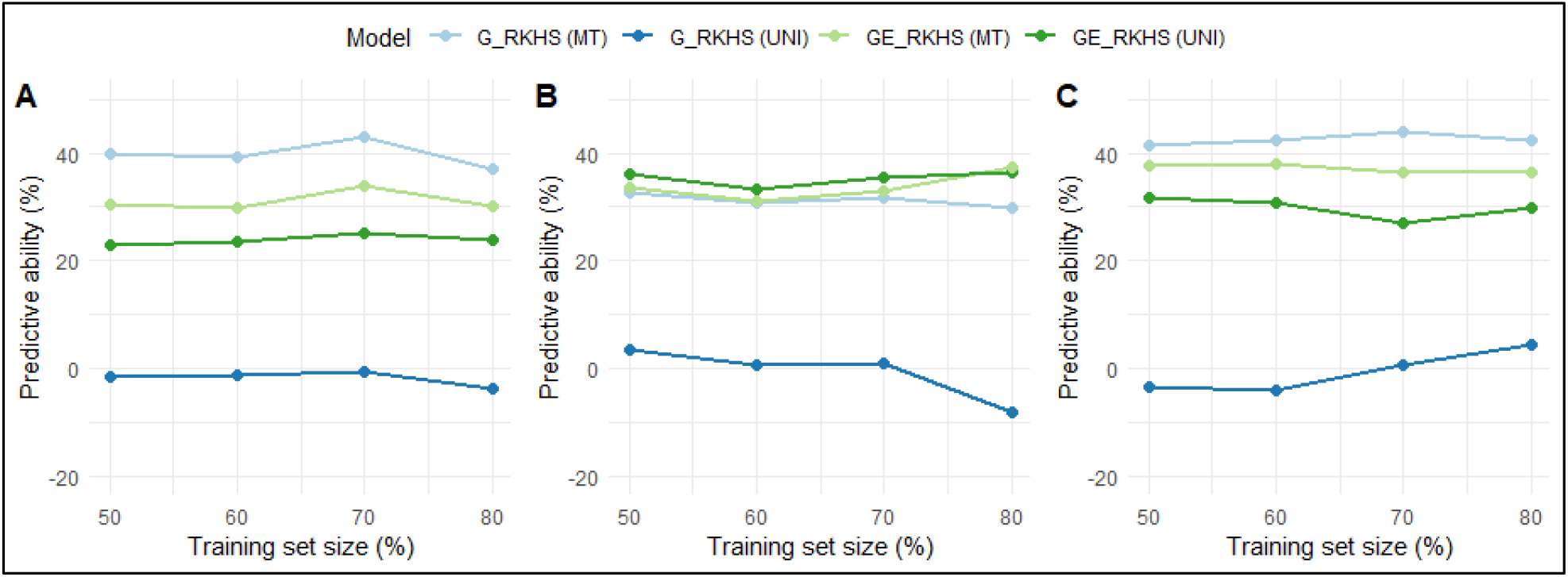
Average predictive abilities with increasing training population set size using RKHS model for seed yield, G is prediction model considering genotype, GE is prediction model integrating GxE interaction, MT is multivariate, UNI is univariate. (A) MOT21 and CAR22 datasets trained to predict MOT20, (B) MOT20 and CAR22 datasets trained to predict MOT21, (C) MOT20 and MOT21 datasets trained to predict CAR22.

**Figure 3.**
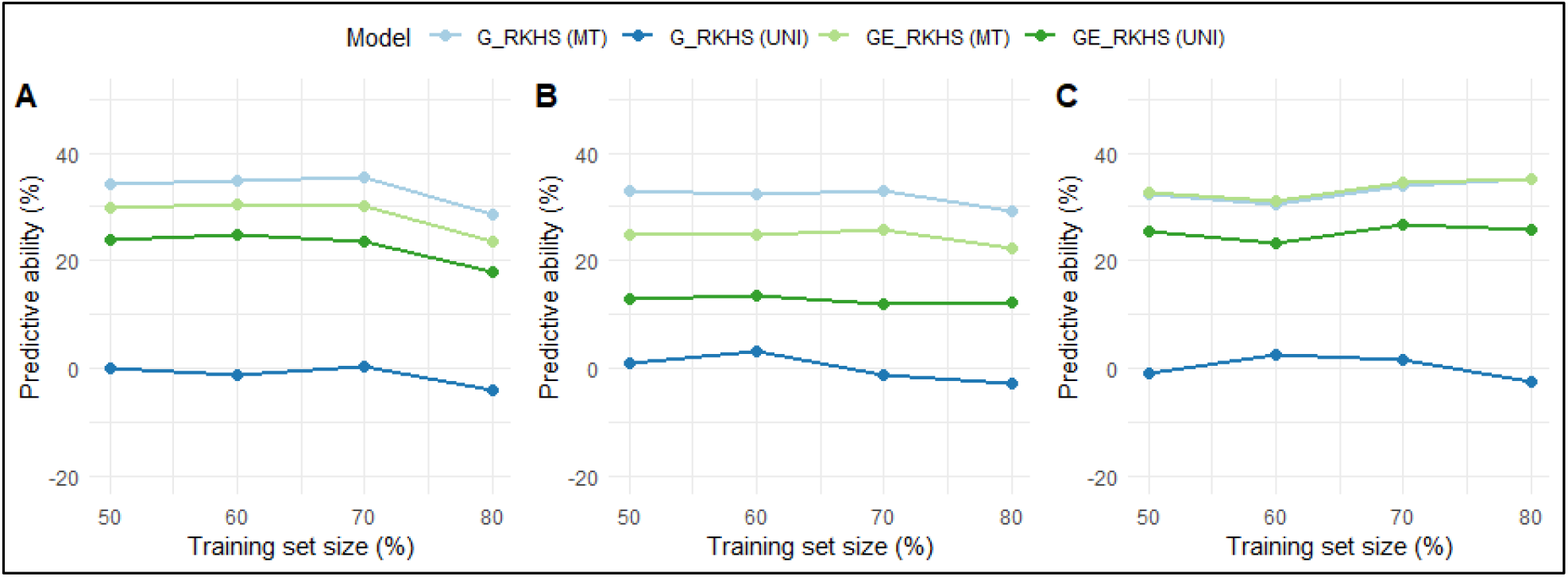
Average predictive abilities with increasing training set size using RKHS model for seed protein content, G is prediction model considering genotype, GE is prediction model integrating GxE interaction, MT is multivariate, UNI is univariate. (A) MOT21 and CAR22 datasets trained to predict MOT20, (B) MOT20 and CAR22 datasets trained to predict MOT21, (C) MOT20 and MOT21 datasets trained to predict CAR22.

## CONCLUSION

Our research findings highlight the intricate dynamics of genomic prediction for seed yield and seed protein content in the face of diverse environmental conditions. The consistent superiority of the RKHS model, particularly in capturing GxE, highlights its robustness and as a choice model in GS. Furthermore, the adoption of MTME-GP has proven instrumental in addressing the complexities associated in predicting inherently low trait heritabilities such as grain yield and total protein content. To fully harness the potential of genomic prediction in plant breeding, composition of the training set in terms of the individuals as well as the heritability of the environments for MTME-enabled GS should be carefully considered. More so, including a wider array of correlated traits in prediction models, integrating deep learning for a more profound understanding of genetic architecture, and incorporating multi-omics data for a comprehensive view of trait genetics and molecular foundations all hold promise. This research marks a significant stride towards unlocking the potential of genomics in public plant breeding programs and offers valuable insights into the challenges and opportunities entailed by complex traits and diverse environments.

## Abbreviations

BLUE: Best linear unbiased estimate
BRR: Bayesian GBLUP model
CAR22: Carrington 2022 dataset
G_RKHS: RKHS model excluding GxE interaction
GE_RKHS: RKHS model accounting for GxE interaction
GEBV: Genomic estimated breeding value
GP: Genomic prediction
GS: Genomic selection
GxE: genotpe-by-environment interaction
MAGIC: multi-parent advanced generation inter-cross
MOT20: Minot 2020 dataset
MOT21: Minot 2021 dataset
MT: Multivariate or Multi-trait
MTME: Multi-trait multi-environment
MTME-GP: Multi-trait multi-environment genomic prediction
NCREC: North Central Research Extension Center
NDSU: North Dakota State University
NIR: Near infrared spectroscopy
PA: Predictive ability
PYT: Preliminary yield trial
RKHS: Reproducing Kernel Hilbert Spaces model
UNI: Univariate

## ACKNOWLEDGMENTS

The authors express gratitude for the funding for this research from the North Dakota Department of Agriculture through the Specialty Crop Block Grant Program (19-429). Additionally, appreciation is extended to Bandillo Lab members for their assistance in conducting field research and collecting data.

## DATA AVAILABILITY STATEMENT

The SNP dataset utilized in this study is accessible through: https://www.ncbi.nlm.nih.gov/sra/PRJNA730349. For access to the phenotype data, please contact the corresponding author.

