## Supplemental figures for "Multi-trait multi-environment genomic prediction of preliminary yield trials in pulse crops"

### SUPPLEMENTAL MATERIAL

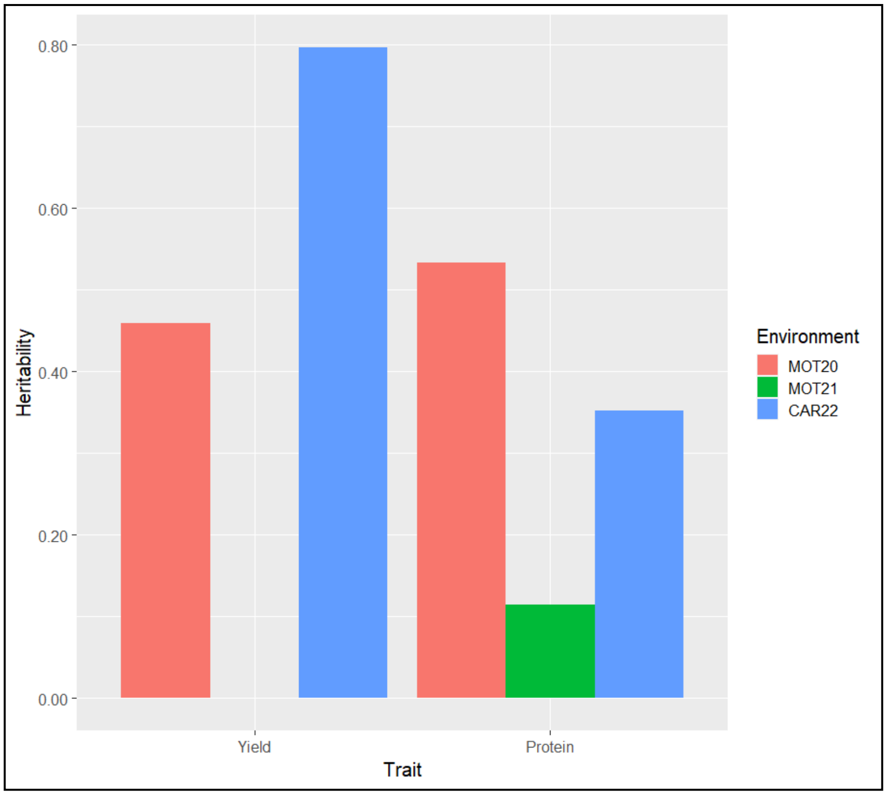


**Supplementary Figure 1. Heritability estimates for yield and protein under three environments, MOT20 is Minot 2020, MOT21 is Minot 2021, CAR22 is Carrington 2022.**

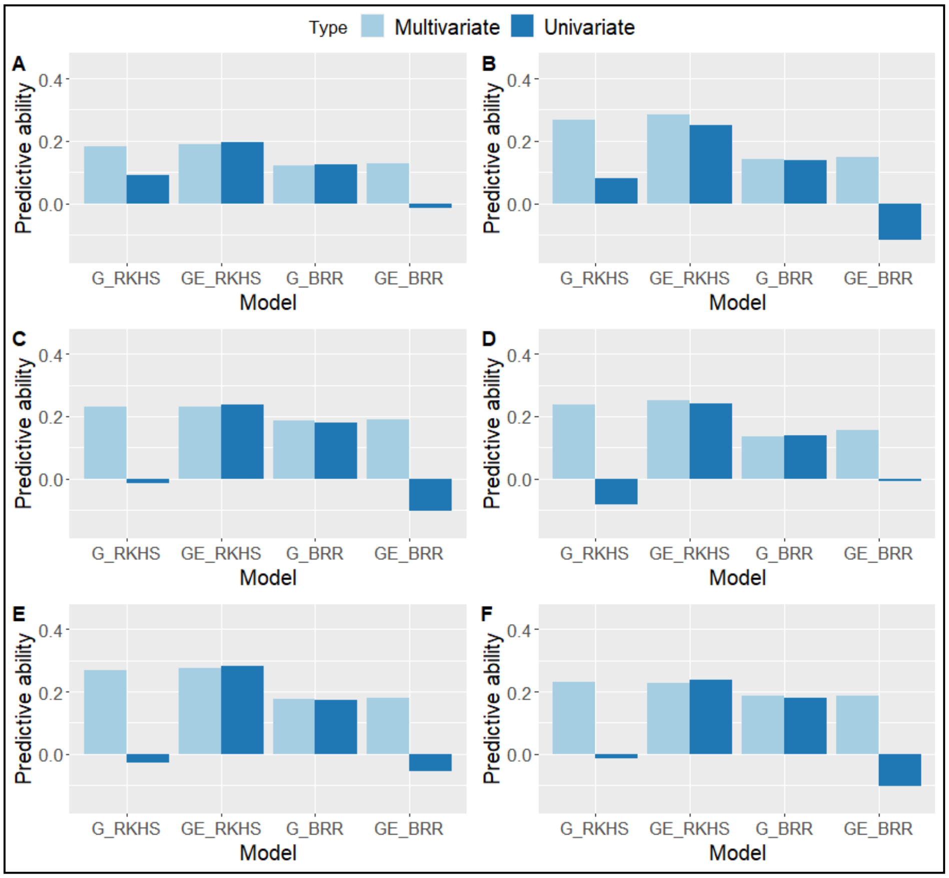


**Supplementary Figure 2: Predictive ability for seed yield using different genomic prediction models across single environments, BRR is Bayesian GBLUP model, RKHS is Reproducing Kernel Hilbert Spaces model, MT is multivariate, UNI is univariate, G is prediction model considering genotype, GE is prediction model integrating GxE interaction. (A) MOT21 dataset trained to predict MOT20, (B) CAR22 dataset trained to predict MOT20, (C) MOT20data set trained to predict MOT21, (D) CAR22 dataset trained to predict MOT21, (E) MOT20 dataset trained to predict CAR22, (F) MOT21 dataset trained to predict CAR22.**

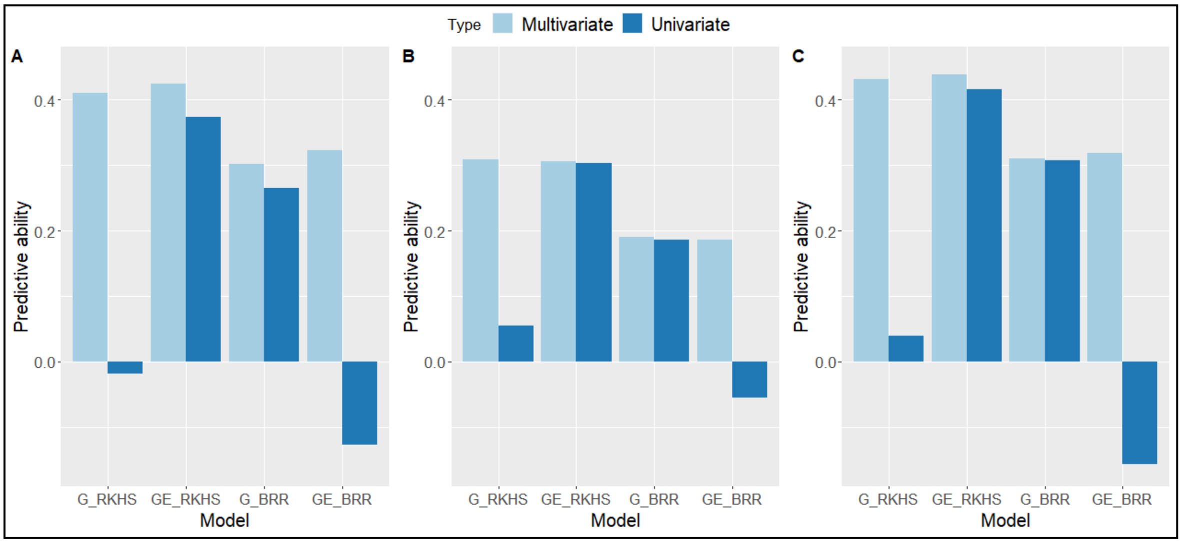


**Supplementary Figure 3: Predictive ability for seed yield using different genomic prediction models across multiple environments, BRR is Bayesian GBLUP model, RKHS is Reproducing Kernel Hilbert Spaces model, MT is multivariate, UNI is univariate, G is prediction model considering genotype, GE is prediction model integrating GxE interaction. (A) MOT21 and CAR22 datasets trained to predict MOT20, (B) MOT20 and CAR22 datasets trained to predict MOT21, (C) MOT20 and MOT21 datasets trained to predict CAR22.**

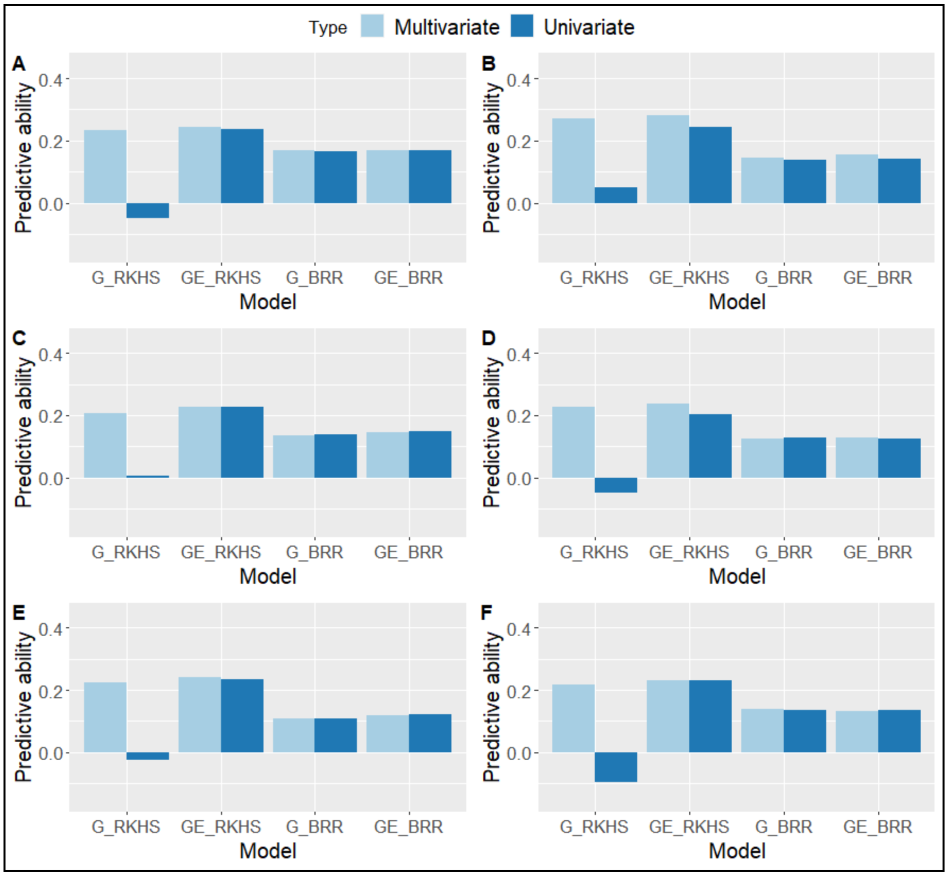


**Supplementary Figure 4: Predictive ability for seed protein content using different genomic prediction models across single environments, BRR is Bayesian GBLUP model, RKHS is Reproducing Kernel Hilbert Spaces model, MT is multivariate, UNI is univariate, G is prediction model considering genotype, GE is prediction model integrating GxE interaction. (A) MOT21 dataset trained to predict MOT20, (B) CAR22 dataset trained to predict MOT20, (C) MOT20data set trained to predict MOT21, (D) CAR22 dataset trained to predict MOT21, (E) MOT20 dataset trained to predict CAR22, (F) MOT21 dataset trained to predict CAR22.**

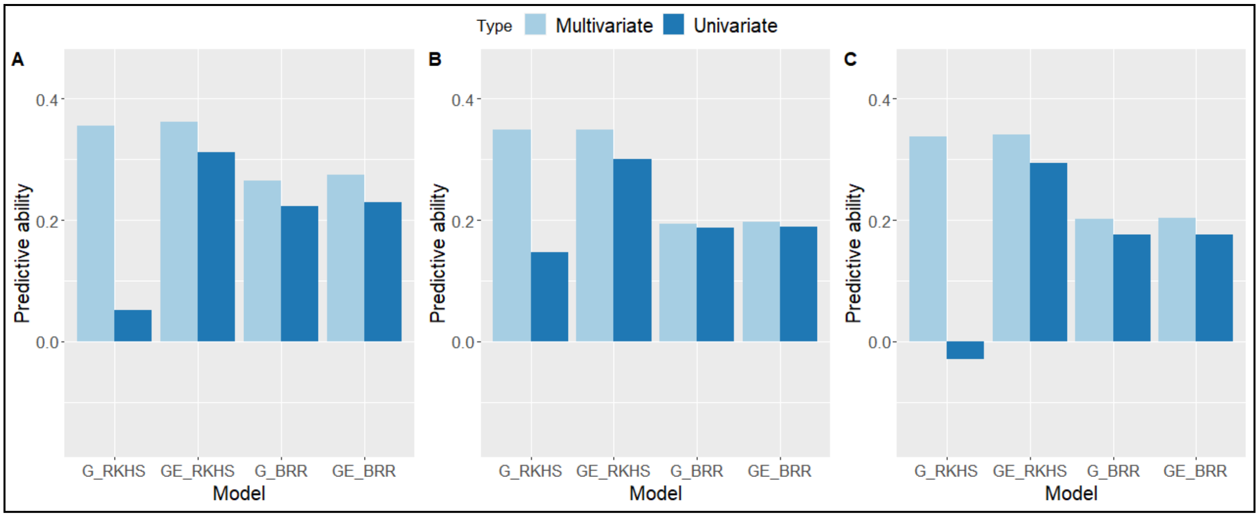


**Supplementary Figure 5: Predictive ability for seed protein content using different genomic prediction models across multiple environments, BRR is Bayesian GBLUP model, RKHS is Reproducing Kernel Hilbert Spaces model, MT is multivariate, UNI is univariate, G is prediction model considering genotype, GE is prediction model integrating GxE interaction. (A) MOT21 and CAR22 datasets trained to predict MOT20, (B) MOT20 and CAR22 datasets trained to predict MOT21, (C) MOT20 and MOT21 datasets trained to predict CAR22.**

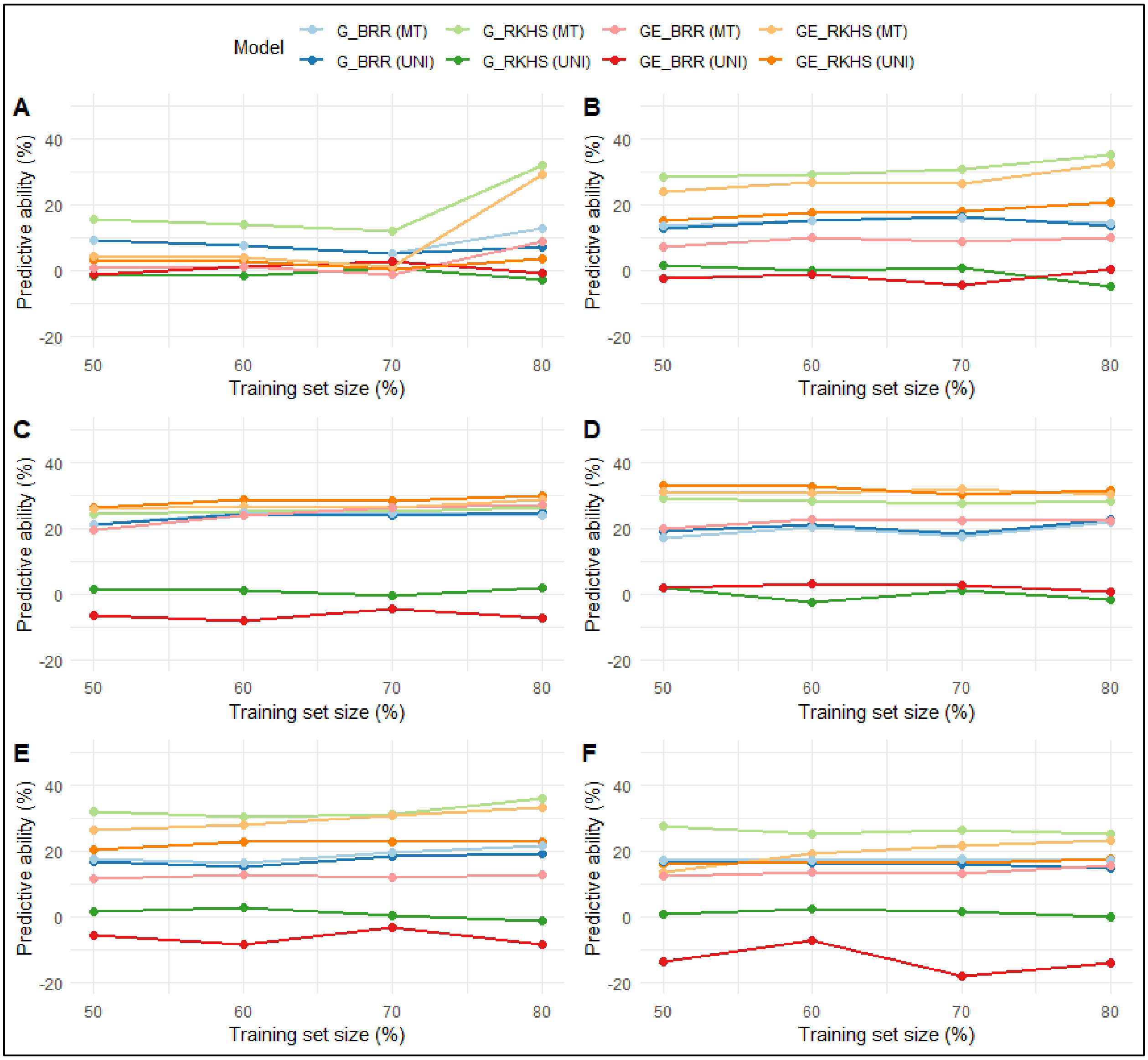


**Supplementary Figure 6: Average predictive abilities with increasing training set size using different genomic prediction models for seed yield, BRR is Bayesian GBLUP model, RKHS is Reproducing Kernel Hilbert Spaces model, MT is multivariate, UNI is univariate, G is prediction model considering genotype, GE is prediction model integrating GxE interaction. (A) MOT21 dataset trained to predict MOT20, (B) CAR22 dataset trained to predict MOT20, (C) MOT20data set trained to predict MOT21, (D) CAR22 dataset trained**

to predict MOT21, (E) MOT20 dataset trained to predict CAR22, (F) MOT21 dataset trained to predict CAR22.

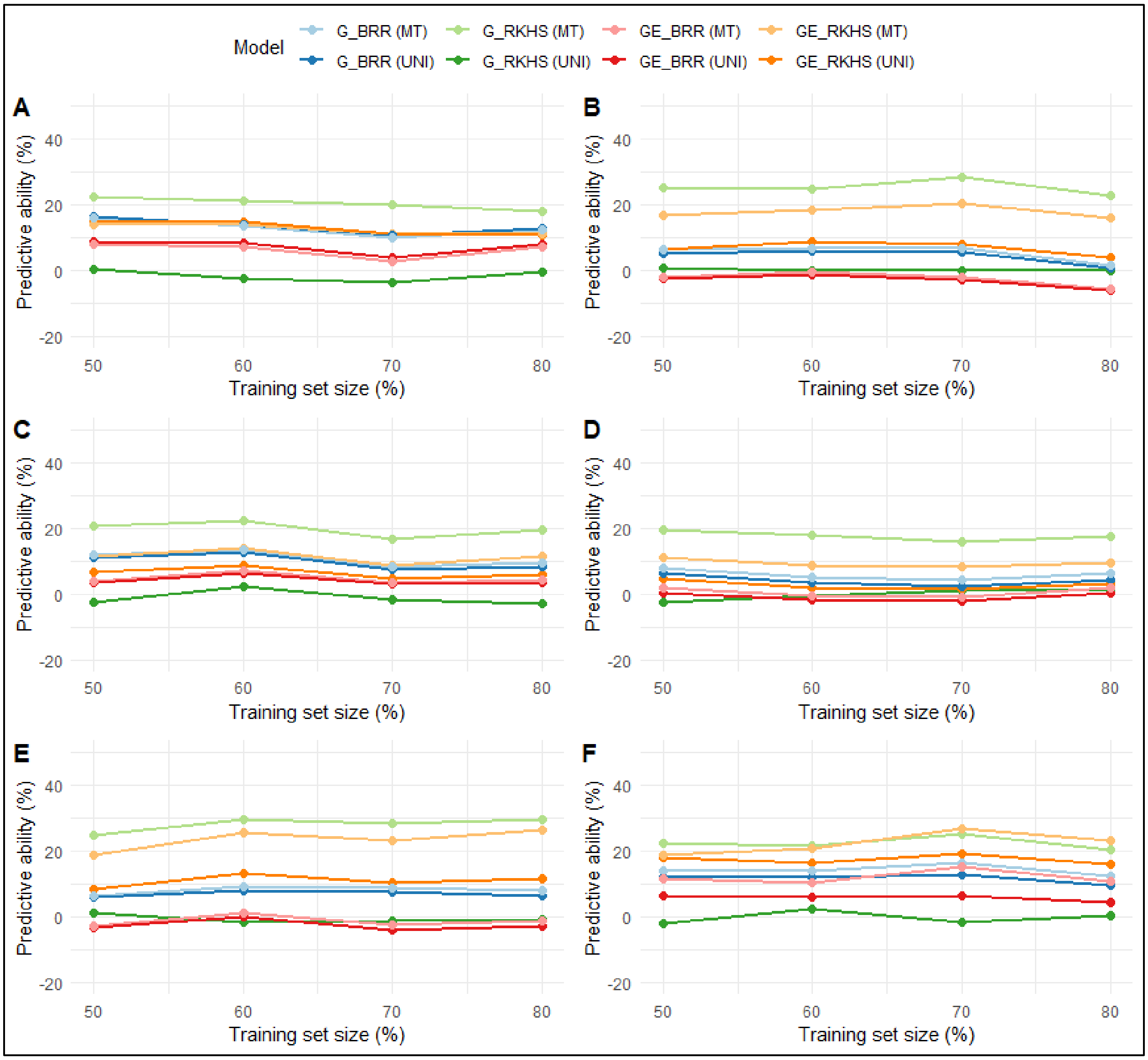


**Supplementary Figure 7: Average predictive abilities for different training set size using different genomic prediction models for seed protein content, BRR is Bayesian GBLUP model, RKHS is Reproducing Kernel Hilbert Spaces model, MT is multivariate, UNI is univariate, G is prediction model considering genotype, GE is prediction model integrating**

**GxE interaction. (A) MOT21 dataset trained to predict MOT20, (B) CAR22 dataset trained to predict MOT20, (C) MOT20data set trained to predict MOT21, (D) CAR22 dataset trained to predict MOT21, (E) MOT20 dataset trained to predict CAR22, (F) MOT21 dataset trained to predict CAR22.**

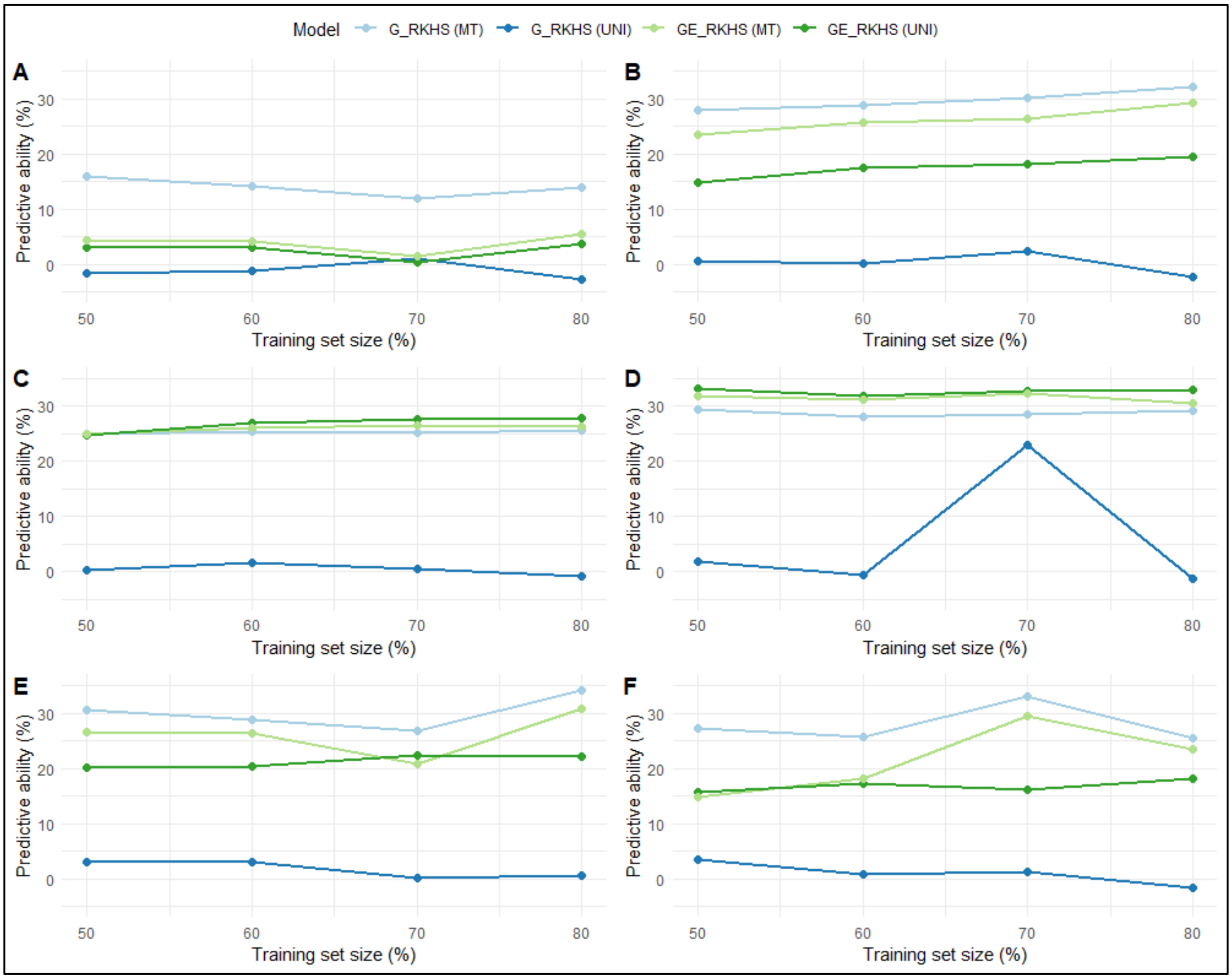


**Supplementary Figure 8: Predictive ability with increasing training population size using RKHS model for seed yield, RKHS is Reproducing Kernel Hilbert Spaces, MT is multivariate, UNI is univariate, G is prediction model considering genotype, GE is prediction model integrating GxE interaction. (A) MOT21 dataset trained to predict**

**MOT20, (B) CAR22 dataset trained to predict MOT20, (C) MOT20data set trained to predict MOT21, (D) CAR22 dataset trained to predict MOT21, (E) MOT20 dataset trained to predict CAR22, (F) MOT21 dataset trained to predict CAR22.**
